# Toxicity of the model protein 3×GFP arises from degradation overload, not from aggregate formation

**DOI:** 10.1101/2024.02.14.580286

**Authors:** Shotaro Namba, Hisao Moriya

## Abstract

While protein aggregation can cause cytotoxicity, it also forms to mitigate cytotoxicity from misfolded proteins, though the nature of these contrasting aggregates remains unclear. We previously found that overproduction (op) of a three green fluorescent protein linked protein (3×GFP) in yeast cells induces giant aggregates, and is detrimental to growth. Here, we investigated the mechanism of growth inhibition by 3×GFP-op using non-aggregative 3×MOX-op as a control. The 3×GFP aggregates were induced by misfolding, and 3×GFP-op had higher cytotoxicity than 3×MOX-op because it perturbs the ubiquitin-proteasome system. Static aggregates formed by 3×GFP-op dynamically trapped Hsp70, causing the heat shock response. Systematic analysis of mutants deficient in the protein quality control suggested that 3×GFP-op did not cause critical Hsp70 depletion and that the formation of aggregates functioned in the direction of mitigating toxicity. Artificial trapping of essential cell cycle regulators into 3×GFP aggregates caused abnormalities in the cell cycle. In conclusion, the formation of the giant 3×GFP aggregates itself is not cytotoxic, as it does not entrap and deplete essential proteins. Rather, it is productive, inducing the heat shock response while preventing an overload to the degradation system.

## Introduction

Most proteins need to fold into the correct three-dimensional structures to perform their functions properly (Dobson, Šali, and Karplus 1998). However, mutations, translation errors, high temperatures, and oxidative stress can affect the protein folding process and induce misfolding. Misfolded proteins are not only useless but sometimes form cytotoxic aggregates (Dill and MacCallum 2012; Lindquist and Kelly 2011). These aggregates are associated with cellular dysfunction due to high temperature and oxidative stress, as well as aging and neurodegenerative diseases (Aguzzi and Lakkaraju 2016; Sweeney et al. 2017). Thus, there are intracellular protein quality control systems, proteostasis, to prevent protein misfolding and to deal with aggregates (Hartl, Bracher, and Hayer-Hartl 2011; Sinnige, Yu, and Morimoto 2020).

Aggregates have long been considered cytotoxic, but it is now clear that aggregates are not immediately cytotoxic, but can be toxic in a variety of different ways depending on their size, structure, and composition (Iadanza et al. 2018; Bucciantini et al. 2002). In recent years, it is also believed that aggregates are actively formed as a quality control mechanism to isolate cytotoxic misfolded proteins (Rothe, Prakash, and Tyedmers 2018; Kaganovich, Kopito, and Frydman 2008). The nature of protein aggregates has been investigated for disease-related proteins, particularly using budding yeast as a model eukaryotic cell. Polyglutamine (PolyQ), linked to Huntington’s disease, forms amorphous/mesh-like aggregates or amyloid fibrils, leading to ER and heat stress responses (Klaips et al. 2020). Similarly, intracellular aggregates of α-synuclein, associated with Parkinson’s disease, disrupt transport between the ER and Golgi apparatus (Cooper et al. 2006; Lázaro et al. 2014). Studies of VHL, UBC9-TS, and TDP-43 showed that toxicity is caused by highly reactive misfolded proteins and very small aggregates (Kaganovich, Kopito, and Frydman 2008; Bolognesi et al. 2019). Interestingly, aggregates of polyQ protein, which normally have an amyloid-like structure, become a mesh structure capable of dynamically trapping Hsp70 upon overexpression of the cochaperone Ydj1 (Klaips et al. 2020). In this state, a productive heat shock response can be triggered. Thus, the aggregates themselves can be cytoprotective. Furthermore, cells have mechanisms to sequester toxic misfolded proteins into intracellular compartments called JUNQ and IPOD, which are observed as large aggregates (Miller et al. 2015; Hill et al. 2016). In contrast to primary quality control, such as avoidance of protein misfolding and rapid degradation of misfolded proteins, the mechanism to isolate such misfolded proteins is called spatial protein quality control (Hill, Hanzén, and Nyström 2017; Rothe, Prakash, and Tyedmers 2018).

We recently found that overproduction (op) of EGFP, a widely used green fluorescent protein in yeast cells promotes a heat shock response, presumably by misfolding, and the formation of aggregates containing Hsp70 (Namba et al. 2022). On the other hand, moxGFP (MOX), which has improved folding properties and lacks cysteine, did not produce such aggregates. Furthermore, the overproduction of three EGFP-linked proteins (3×GFP-op) causes stronger growth inhibition (more toxic) than EGFP-op (Kintaka et al. 2020). In 10-20% of 3×GFP-op cells, one giant aggregate, about 5 μm in size, with GFP fluorescence is produced. In the insoluble fraction of 3×GFP-op cells, the molecular chaperone Hsp70 (Ssa1/Ssa2) and the glycolytic enzymes Fba1 and Eno2 are enriched, suggesting that the aggregates may be toxic by sequestering these proteins. On the other hand, the ubiquitinated 3×GFP accumulates in the same insoluble fraction, and 3×GFP-op has a negative genetic interaction with proteasome mutants. This also suggests that the overload on degradation caused by misfolded 3×GFP is responsible for growth inhibition. Considering the above, the 3×GFP-op cells were analyzed in more detail in this study, as they are a good model case to investigate the cytotoxicity induced by protein misfolds and aggregates.

One limitation in the study of whether aggregation is a source of cytotoxicity or a mechanism for avoiding toxicity, and how it is accomplished, is often the lack of appropriate control proteins (Schneider, Nyström, and Widlund 2018). Since the massive expression of a protein is itself a source of cytotoxicity for a variety of reasons (Moriya 2015), the formation of aggregates itself must be distinguishable from the cytotoxicity produced by an excess of that protein. In this study, 3×MOX-op was used as a control because the molecular weight and structure of 3×GFP and 3×MOX are close, which should allow us to better discriminate between the effects of aggregation and other effects, such as the effect of high expression of 75 kD proteins itself. We investigated the conditions for the generation of giant aggregates of 3×GFP, the components they contain, the formation process, and the mechanism by which they exert cytotoxicity while using 3×MOX as a control. We show that 3×GFP-op creates large aggregates that trap Hsp70 but are not themselves the cause of toxicity, but rather an overload of degradation by the ubiquitin-proteasome system. We also show that the entrapment of essential proteins into aggregates can be a potential mechanism of toxicity.

## Results

### 3×MOX can be a non-aggregating control for 3×GFP

To separate the effects of high-level protein expression from those caused by aggregation, we first generated 3×MOX, a control protein with a similar structure to 3×GFP but without aggregation potential. We overproduced 3×GFP and 3×MOX under the constitutive *PYK1* promoter (*PYK1_pro_*) or inducible *WTC_846_* system (Azizoglu, Brent, and Rudolf 2021) on the pTOW40836 plasmid (Makanae et al. 2013)(Figure 1A). This plasmid is a multicopy plasmid whose copy number increases under SC–LU medium condition up to >100 copies per cell (Moriya, Shimizu-Yoshida, and Kitano 2006; Makanae et al. 2013). We confirmed the development of single large aggregates in the 3×GFP-op cells as previously reported (Kintaka et al. 2020), while there was no aggregate in 3×MOX-op cells (Figure 1B). Conditions triggering protein misfolding such as 5-azacytidine (AZC: mistranslation) and hydrogen peroxide (H_2_O_2_: oxidative stress) treatment enhanced the aggregate formation upon 3×GFP-op, but not 3×MOX-op (Figure 1C-E). In the aggregate, 3×GFP should not be completely but partially misfolded because GFP fluorescence is still observed even when aggregated (Figure 1B). These results suggest that the aggregates are formed with partially misfolded GFP, but not by the clustering of folded GFPs, and 3×MOX can be used as a control non-aggregative protein for 3×GFP.

**Figure 1.**
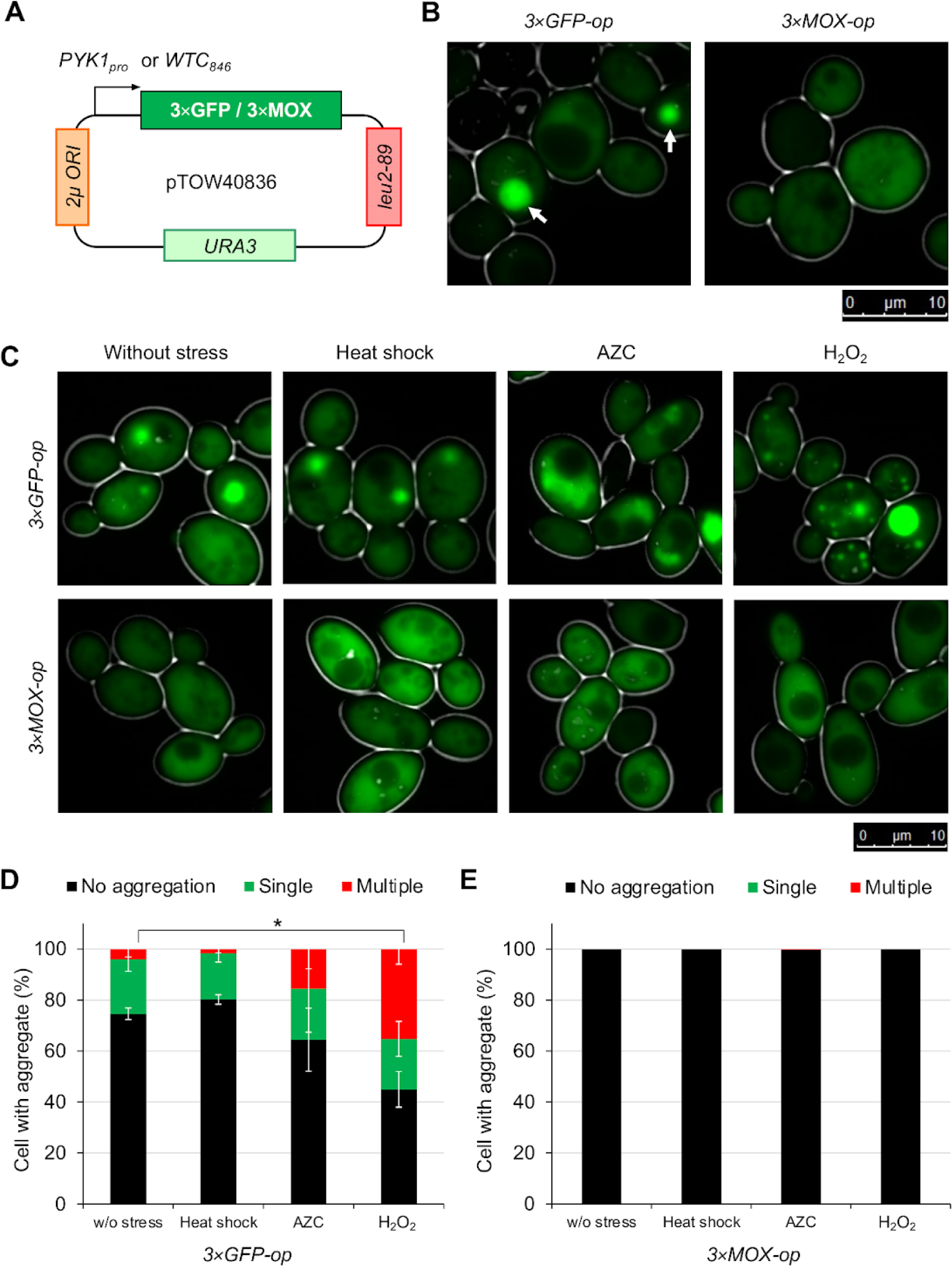
3×MOX can be a non-aggregating control for 3×GFP. **(A)** Plasmids used in this study. 3×GFP and 3×MOX were overproduced under the *PYK1* promoter (*PYK1_pro_*) or *WTC_846_* on the multicopy plasmid pTOW40836. The nucleotide sequences of the three linked GFPs and MOXs are different to prevent accidental homologous recombination. **(B)** Fluorescence microscopy images of the cells under 3×GFP-op and 3×MOX-op. Representative images of log-phase cells cultured in SC–LU medium are shown. The arrowheads indicate aggregates. **(C)** Fluorescence microscopic images of cells under indicated stress conditions under 3×GFP-op and 3×MOX-op. Representative images are shown. **(D and E)** Quantification of aggregation of 3×GFP-op (**D**) and 3×MOX-op (**E**) cells under indicate stress conditions. *: *p* < 0.05; n = 3; using the Dunnett’s method.

### 3×GFP has stronger cytotoxicity than 3×MOX

We next compared the cytotoxicity of 3×GFP and 3×MOX. We also analyzed monomeric MOX to assess the increased cytotoxicity resulting from the linking of three MOX. We first measured the maximum growth rate upon a stepwise increase in the expression using the *WTC_846_* system (Figure 2A). We note that even without the inducer anhydrous tetracycline (aTc), 3×GFP-op and 3×MOX-op cells showed reduced growth rate compared to the vector control under the high-copy conditions (SC–LU, in Figure 2A). This suggested that the leaked expression of these proteins already caused some cytotoxicity. Upon induction up to 150 nM aTc, all three proteins showed gradual decreases in the growth rates. With the aTc concentration above 100 nM, 3×GFP-op cells almost completely halted growth, while 3×MOX-op and MOX-op cells maintained growth. In addition, the growth of MOX-op cells was always better than that of 3×MOX-op cells under all conditions, suggesting that cytotoxicity was higher for 3×GFP, 3×MOX, and MOX, in that order.

We next assessed the maximum protein levels at which the cells could maintain growth (expression limit). We analyzed the protein levels of 3×GFP at 50 nM aTc, 3×MOX at 50 nM, 250 nM, and 500 nM aTc (Figure 2B). We then measured the expression limits as a % of the MOX limit (expressed under *TDH3pro* which is equivalent to the maximum induction of the *WTC_846_*system) (Figure S1). The maximum measurable expression limit of 3×GFP (at 50 nM aTc; 9 unit) was about 1/3 of that of 3×MOX (500 nM aTc; 38 unit), suggesting that 3×GFP has stronger cytotoxicity than 3×MOX. We note that the linking of three MOX itself increased cytotoxicity because the expression limit of 3×MOX is lower than that of MOX (Figure 2B).

**Figure 2.**
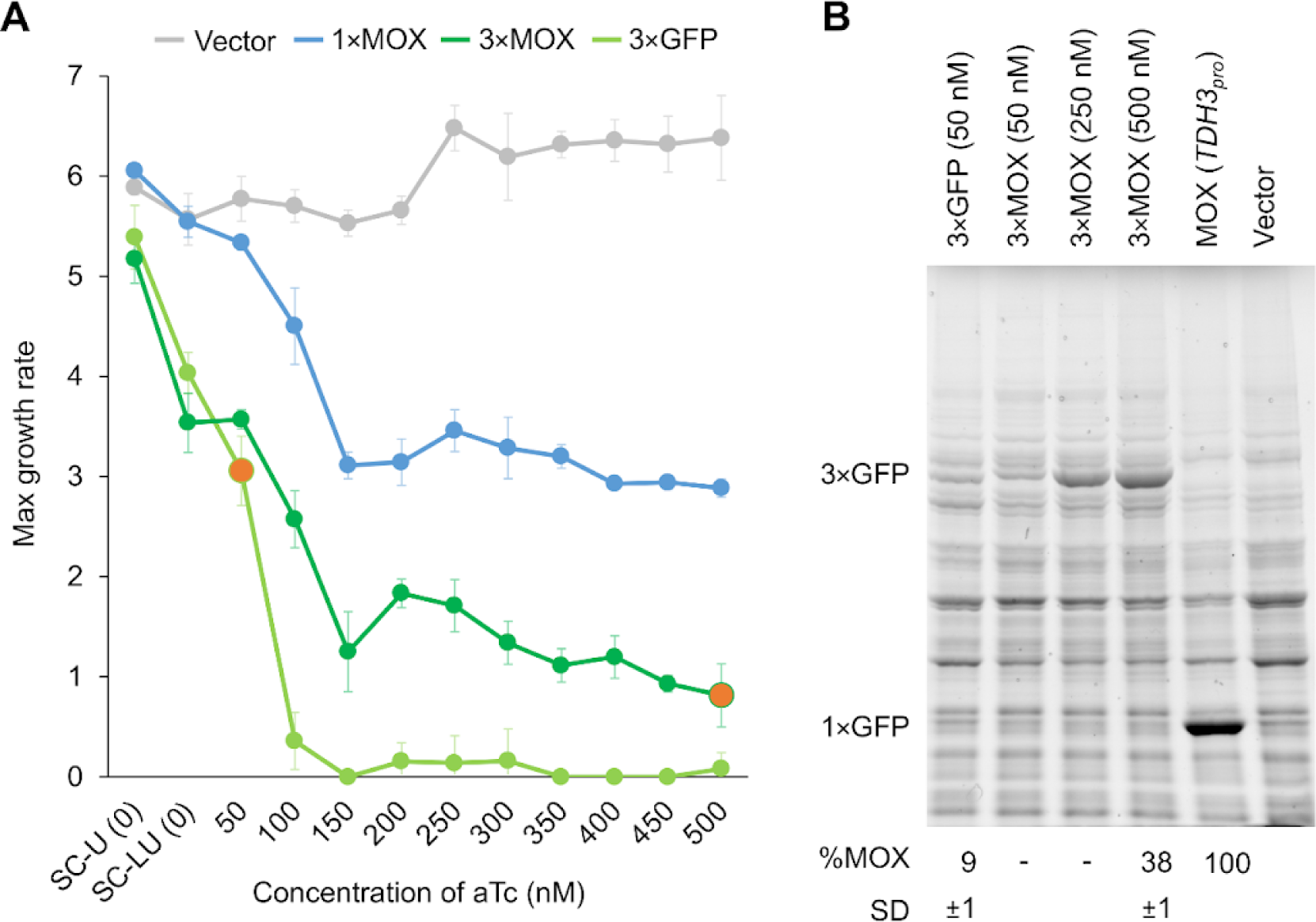
3×GFP has stronger cytotoxicity than 3×MOX. **(A)** Max growth rates of cells at different expression levels. Each protein was expressed under *WTC_846_*on pTOW40836 (Figure 1A). Orange circles indicate the concentrations used for SDS-PAGE measurements in **B**. Anhydrotetracycline (aTc) was not added for SC–U (0) and SC–LU (0). Each point represents the mean of the max growth rate, with error bars representing its standard deviation (n = 4). **(B)** Measurement of expression limits. A gel image of SDS-PAGE of total protein from cells expressing the indicated proteins is shown. The amount of protein corresponding to the size of overproduced protein (shown as 1**×**GFP and 3**×**GFP) was calculated using MOX as 100%, and the SD of the three replicates is also shown (numbers under the gel image). The details of protein quantification are described in Figure S1.

### 3×GFP-op perturbs the ubiquitin-proteasome system

We previously found that 3×GFP-op had negative genetic interactions with proteasome mutants and high-molecular-weight ubiquitinated proteins were accumulated in the insoluble fraction upon 3×GFP-op (Kintaka et al. 2020). These results suggested that 3×GFP-op perturbs the ubiquitin-proteasome system. Here, we further analyzed the genetic interactions with 3×MOX as a control. As shown in Figure 3A, the growth rate of temperature-sensitive mutant strains of three proteasome genes (*pre6-5001*, *pup1-1*, and *rpn12-1*) under 3×GFP-op showed significantly slower growth than of the vector control or under 3×MOX-op. We also tried to confirm the ubiquitination of isolated 3×GFP aggregates (Figure 3B). As shown in Figures 3C and 3D, Ni-carrier purified aggregates from 3×GFP-op cells contained high molecular weight proteins detected by anti-GFP and anti-ubiquitin antibodies, but not 3×MOX-op cells. These results confirmed that 3×GFP-op perturbs the ubiquitin-proteasome system, probably through the massive degradation of the 3×GFP aggregates.

**Figure 3.**
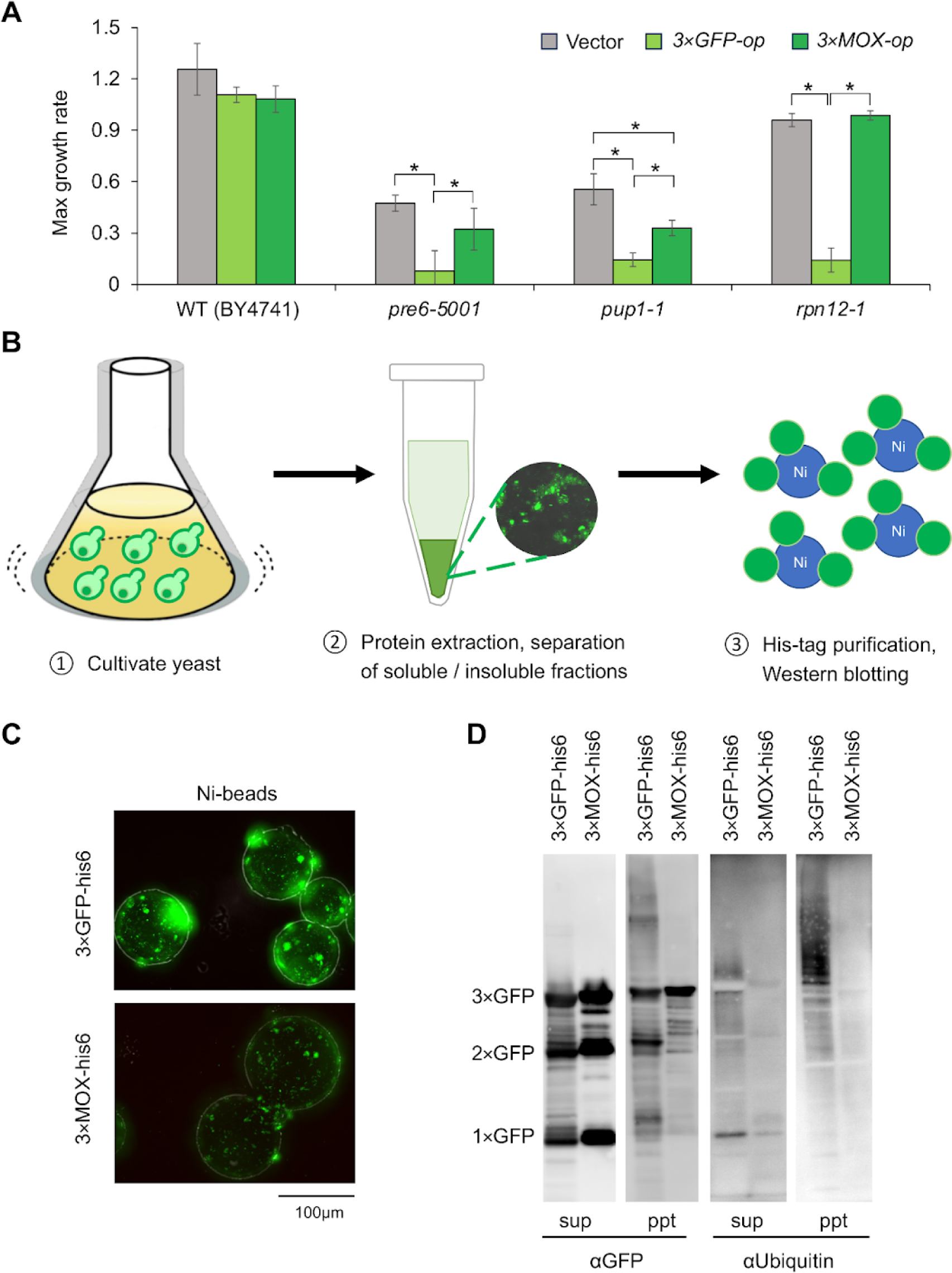
3×GFP-op perturbs the ubiquitin-proteasome system. **(A)** Max growth rates of proteasome mutants under 3×GFP-op, 3×MOX-op, and with the empty vector (Vector). Cells were cultivated at 26°C. The bars represent the means, and the error bars represent the standard deviations (n = 4). **(B)** Schematic diagram of 3×GFP aggregate purification. **(C)** Fluorescence microscopic images of Ni-carrier bound to aggregates isolated from insoluble fractions of cells overproducing 3×GFP-his6 and 3×MOX-his6. **(D)** Western blot analysis of purified 3×GFP-his6 (3×GFP) and 3×MOX-his6 (3×MOX) from soluble (sup) and insoluble (ppt) fractions. GFP and MOX were detected using anti-GFP antibodies (αGFP), and ubiquitinated proteins were detected using anti-ubiquitin antibodies (αUbiquitin). Predicted sizes of 1×GFP, 2×GFP, and 3×xGFP are shown.

### 3×GFP aggregates entrap dynamic Hsp70, which induces a heat shock response

We previously found that Hsp70 (Ssa1/Ssa2) and the glycolytic enzymes Eno2 and Fba1 were enriched in the insoluble fraction of 3×GFP-op cells using LC-MS/MS (Kintaka et al. 2020). To confirm whether these three proteins were included in the aggregation of 3×GFP, here we used fluorescence microscopy. Among these three proteins with a red fluorescent protein (mScarlet-I: mSca) bound to the C-terminus, co-localization of Ssa1-mSca with 3×GFP aggregates was confirmed (Figure 4A and S2). On the other hand, Eno2-mSca and Fba1-mSca did not co-localize with 3×GFP aggregates. As Eno2 and Fba1 are known to fractionate into insoluble fractions under conditions that cause proteotoxicity (Geiler-Samerotte et al. 2011; Weids et al. 2016), these proteins might become insoluble due to the proteotoxic stress produced by 3×GFP-op.

Kpaips et al. have shown that the kinetics of Hsp70 in aggregates distinguished the aggregate properties (Klaips et al. 2020). We thus analyzed the dynamic properties of Ssa1-mSca co-localized with 3×GFP aggregates by fluorescence recovery after photobleaching (FRAP) analysis (Figure 4B). The fluorescence recovery of 3×GFP was only about 10% even after 60 seconds of bleaching, while the fluorescence of Ssa1-mSca recovered immediately after bleaching, reaching 50% in 40 seconds (Figure 4C). This result suggests that 3×GFPs do not form dense aggregates, but rather create a mesh-like structure in which accumulated Hsp70 can dynamically move.

**Figure 4.**
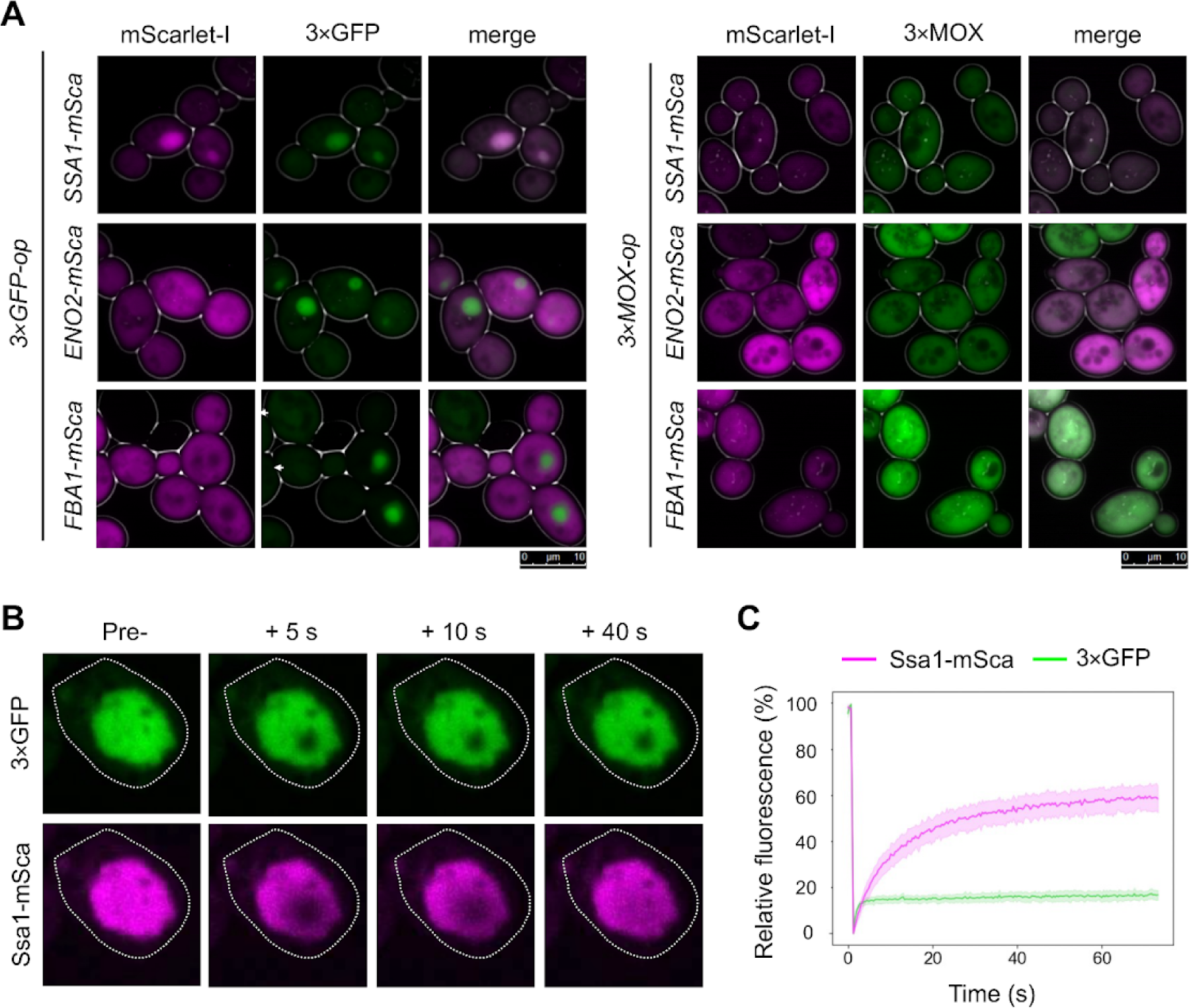
Hsp70 (Ssa1) co-localizes with 3×GFP aggregates, showing dynamic characteristics. **(A)** Fluorescence microscopic images of cells under 3×GFP-op (left) and 3×MOX-op (right) with Ssa1, Eno2, and Fba1 fused with mScarlet-I (mSca). Representative images of mSca-fusion proteins (mScarlet-I), 3×GFP, and 3×MOX, with their merges are shown. **(B)** Fluorescence recovery after photobleaching (FRAP) analysis of GFP and Ssa1-mSca in the 3×GFP aggregates. Shown are representative images in a time series of breaching to a location with a 3×GFP aggregate, where Ssa1-mSca is also co-localized. In all images, the cell shapes are outlined by dotted lines. Pre-: before bleaching; +: seconds (s) after bleaching. White dotted lines indicate the contours of the cells. **(C)** Quantification of FRAP analysis. Relative fluorescence is calculated as 100% before bleaching and 0% immediately after bleaching. The means (solid line) and standard errors (traces) of the 10 cells measured are shown.

We next investigated the process by which 3×GFP and Hsp70 grow into one large aggregate by time-series observation after induction of 3×GFP expression (Figure 5A and B). The 3×GFP was observed as multiple small bright spots at 1 hour after induction. The dots were initially co-localized with Hsp70, but eventually, Hsp70 surrounded the exterior (after 3 hours), and they clustered into a small number of dots (after 6 hours). After a long time, Hsp70 and 3×GFP were mixed into one large aggregate (Overnight). As the aggregates clustered and became huge, the expression of Hsp70 also increased (Figure 5C). Similarly expressed 3×MOX did not induce aggregates (Figure S3).

Combined with the results of the FRAP analysis above, a possible mechanism is that the initial 3×GFP aggregates are gradually dissolved by the surrounding Hsp70 and fuse into a large mesh-like aggregate containing dynamic Hsp70 (Figure 5D). This progression is thought to be similar to the formation process of mesh-like aggregates of polyQ protein with dynamic Hsp70 (Klaips et al. 2020). This polyQ protein aggregate triggers a productive heat shock response by trapping Hsp70. Indeed, 6 hours after induction of 3×GFP expression, Ssa1-mSca fluorescence increased (Figure 5C), suggesting that a heat shock response (HSR) was triggered. Induction of HSR upon 3×GFP-op (but not 3×MOX) was confirmed by the RNAseq analysis (Figure S4, Supplementary data 1). Thus, the large aggregate formation upon 3×GFP-op itself seemed not toxic but protective to the cell.

**Figure 5.**
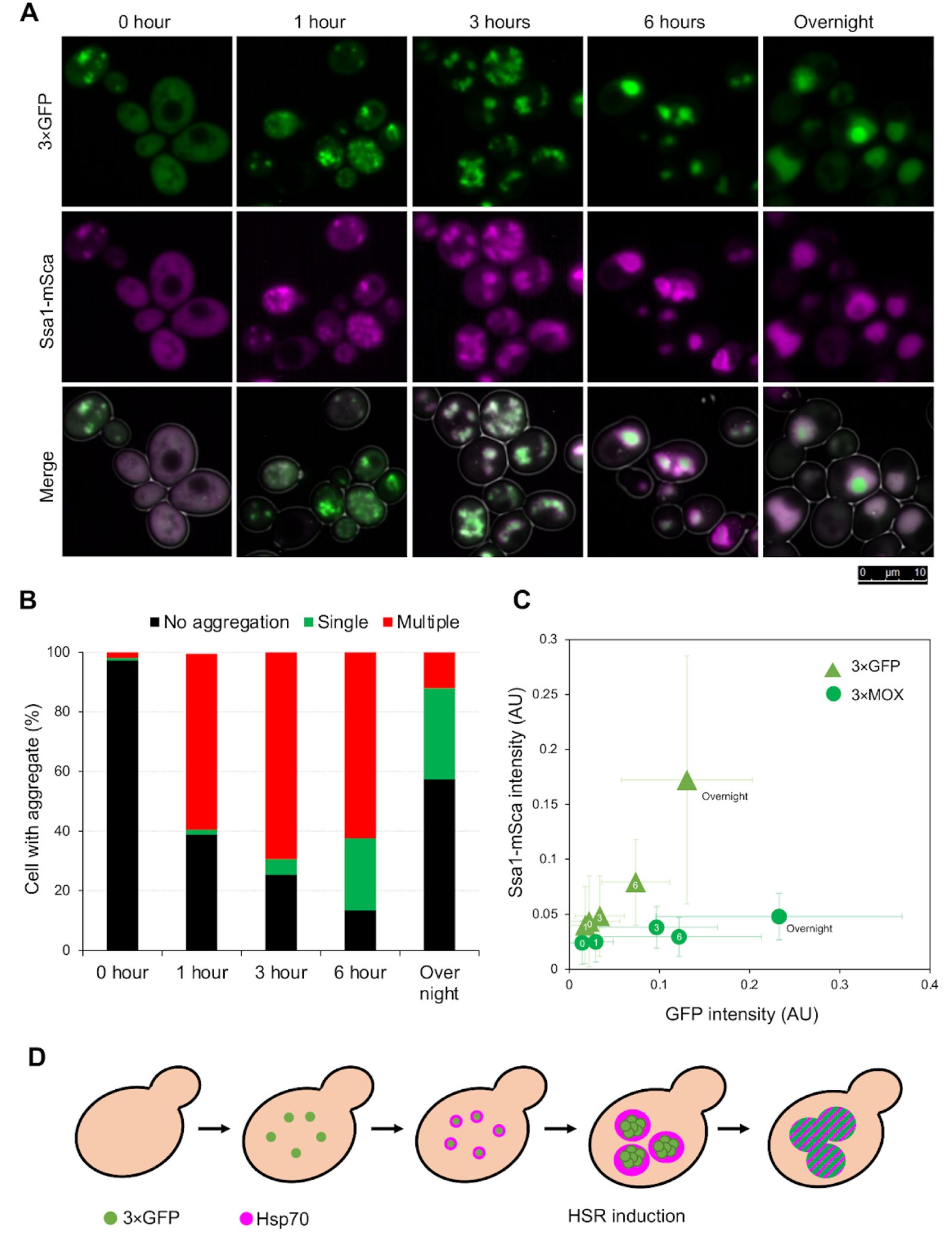
Observation of aggregation process and heat shock response after induction of 3×GFP. **(A)** Time-series images of cells expressing Ssa1-mSca, following the induction of 3×GFP using the *WTC_846_* system (500 nM aTc). Cells at indicated time points were observed. **(B)** Quantification of aggregation of 3×GFP at each time point. At least 100 cells were observed. **(C)** A scatter plot of GFP and Ssa1-mSca intensity following the induction of 3×GFP and 3×MOX. The fluorescence intensities of GFP and mScarlet-I were measured from fluorescence microscopy images of individual cells under induction conditions. The mean values and standard deviations (error bars) from 300 randomly selected cells are shown. The numbers within the markers indicate the time points after induction. **(D)** A model diagram for the 3×GFP aggregation process. The detail is explained in the main text.

### Analysis of secondary quality control mutants suggests that aggregates are protective

To further investigate whether 3×GFP aggregate formation itself is protective, we investigated the relationship between 3×GFP aggregate formation and growth inhibition in 15 gene deletion strains involved in the secondary quality control (Hill, Hanzén, and Nyström 2017)(Figure 6A). We first observed 3×GFP aggregates formation in each deletion strain (Figure 6B, S5A). The deletion strains showed an increase or decrease in the aggregation rates upon 3×GFP-op, while no aggregation was observed in any of the deletion strains upon 3×MOX-op (Figure S5B). We next measured the growth rate of the deletion strains upon 3×GFP-op, 3×MOX-op, and with the vector control (Figure 6C). Again the deletion strains showed an increase or decrease in the growth rates. No significant growth reduction was observed in the *ssa1Δ* and *ssa2Δ* strains compared to the wild type (*p*-value: 0.99 for ssa1Δ and 0.10 for ssa2Δ), suggesting that the trapping of Hsp70 (Ssa1/Ssa2) by the 3×GFP aggregates did not cause growth inhibition caused by Hsp70 depletion. We note that 3×MOX-op mitigates the growth defects of some deletion strains with the vector control. This could be a potential deleterious effect of the experimental system using a high-copy vector (Kintaka et al. 2020).

Next, we examined how an increase or decrease in aggregate formation affects growth (Figure 6D and 6E). The growth rates and aggregation rates in the 3×GFP-op dataset, when the vector was used as a control, showed a weak positive correlation (Pearson’s *r* = 0.33). The correlation was higher (*r* = 0.63) when *ydj1Δ*, which had significantly poorer proliferation, was excluded. Furthermore, the correlation was even higher when 3×MOX-op was used as a control (*r* = 0.79). These results mean that the more aggregates formed, the more growth inhibition was mitigated, suggesting that aggregate formation works in the direction to mitigate cytotoxicity.

**Figure 6.**
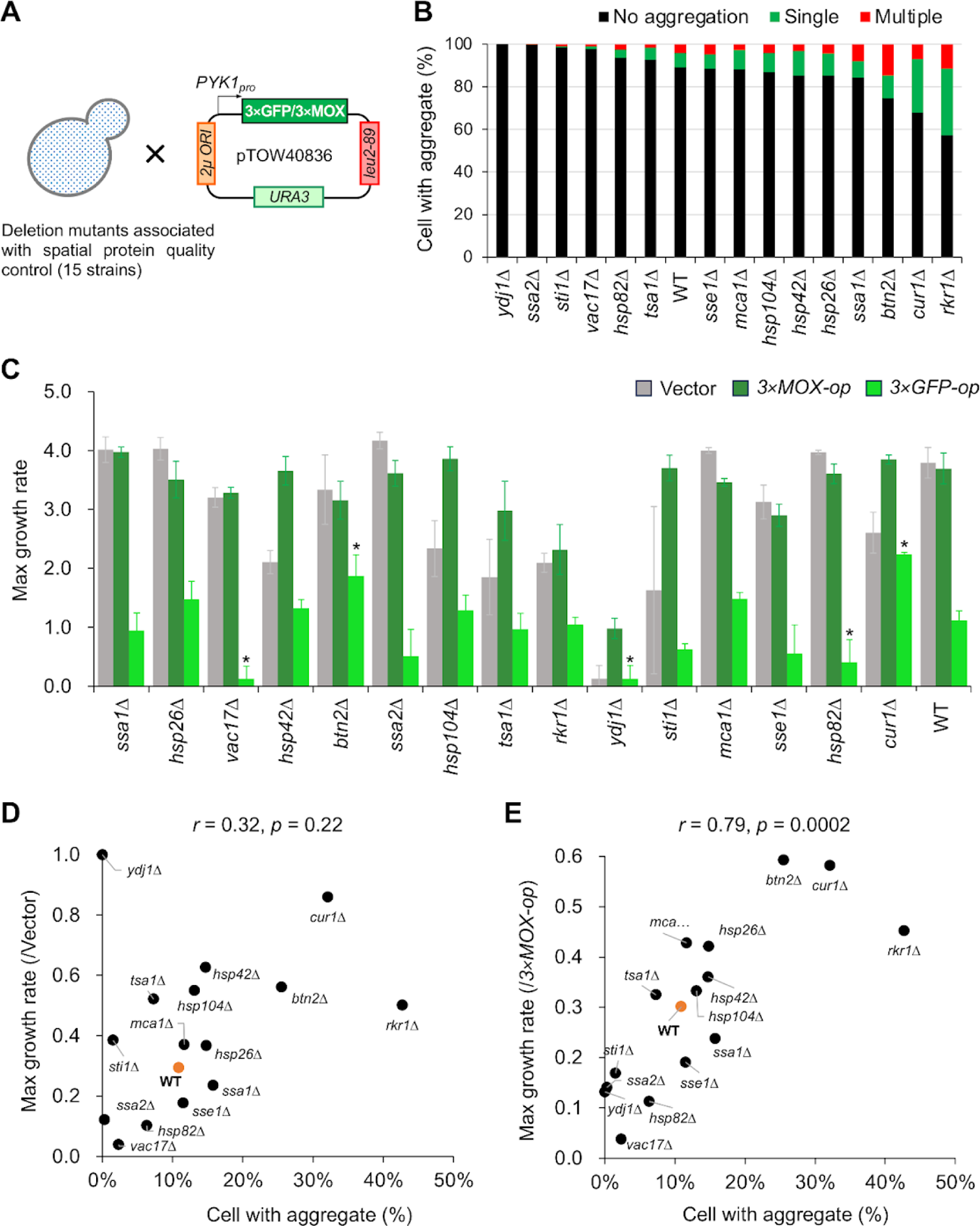
Analysis of secondary quality control mutants suggests that aggregates are protective. **(A)** Schematic presentation of the experiment. **(B)** Quantification of aggregation of mutant cells under 3×GFP-op. At least 100 cells were observed for each mutant strain. **(C)** Max growth rate in each deletion strain under 3×GFP-op, 3×MOX-op, and the vector control (Vector). *: *p* < 0.05; n.s.: *p* >= 0.05; n=3; using Dunnett’s method. **(D and E)** Scatter plots showing the relationships between aggregation propensity and max growth rate across different mutant cells under 3×GFP-op. In **D**, the max growth rate is normalized to the vector control (Vector), while in **E** it is normalized to the 3×MOX-op. *r*: Pearson’s correlation coefficient; *p*: *p*-value of the test for lack of correlation.

### Sequestering essential Cdc proteins into 3×GFP aggregates induces cell cycle dysfunction

The previous results suggest that the cytotoxicity caused by 3×GFP aggregates is not due to sequestration of essential proteins. However, there are known cases in which specific aggregates show cytotoxicity due to the sequestration of essential proteins (Treusch and Lindquist 2012; Park et al. 2013). Therefore, we tested whether these aggregates have the potential to exhibit cytotoxicity when they trap essential proteins.

We accidentally found that monomeric EGFP is incorporated into aggregates when non-fluorescent 3×GFP is overproduced (3×GFP-Y66G) (Figure 7A). We thus considered that co-expression of 3×GFP with GFP-tagged proteins could entrap GFP-tagged proteins into aggregates and deplete them (Figure 7B). To test this idea, 3×GFP was overproduced in strains expressing the GFP-linked Cdc20 and Cdc28, which are essential cell-cycle regulators. As shown in Figures 7C and 7D, cells were highly enlarged upon 3×GFP-op (not 3×MOX-op) in Cdc20-GFP and Cdc28-GFP strains, just as temperature-sensitive mutants of *cdc20* and *cdc28* (Tavormina and Burke 1998; Iida and Yahara 1984). These results indicate that trapping essential proteins into 3×GFP aggregates potentially causes cytotoxicity.

**Figure 7.**
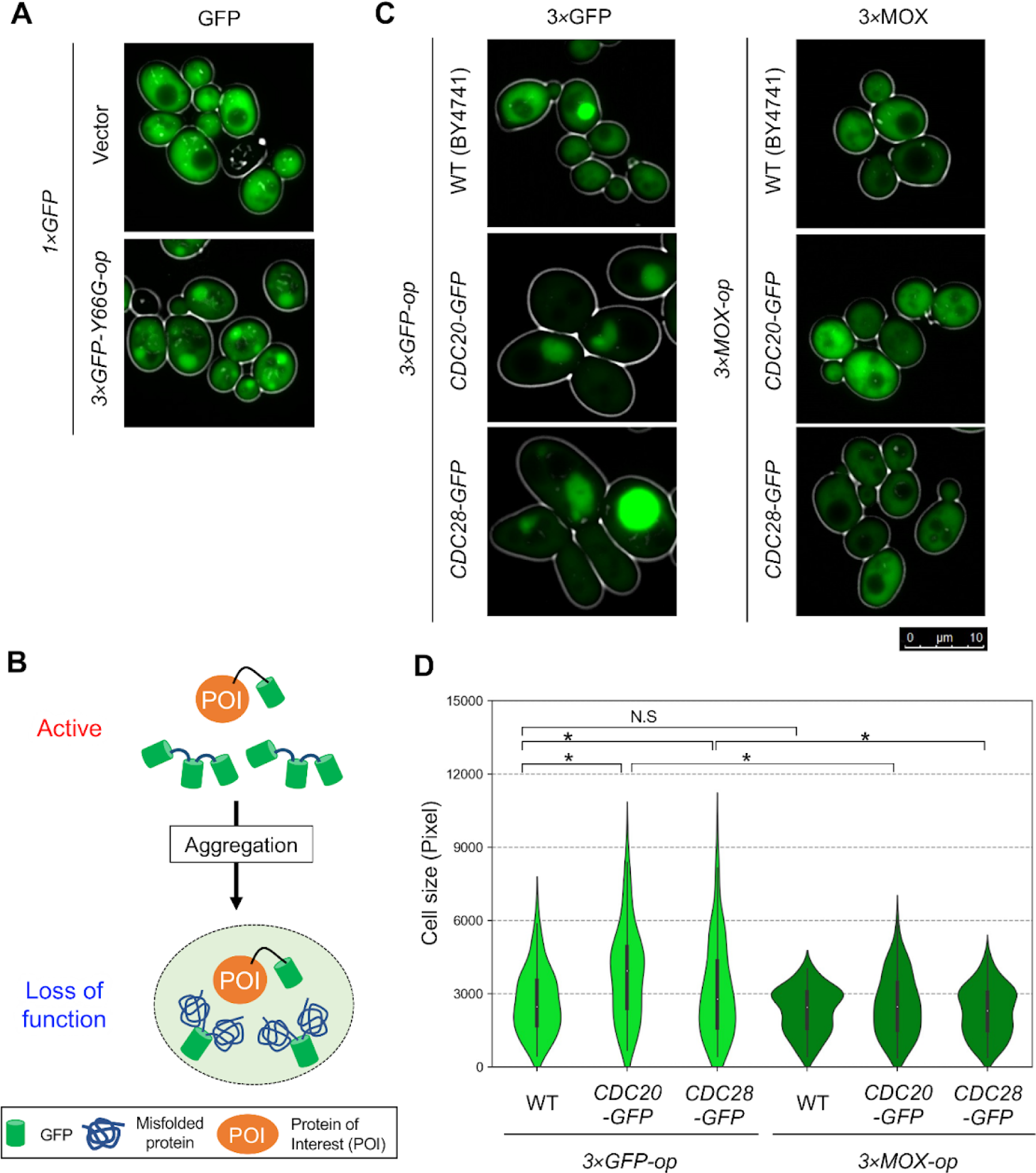
Sequestering essential Cdc proteins into 3×GFP aggregates induces cell cycle dysfunction. **(A)** Fluorescence microscopic images of cells expressing 1×GFP (from *TDH3pro*) with those carrying the vector control (Vector) and cells under 3×GFP-Y66G-op. **(B)** A model diagram showing protein knockdown using aggregates. Normally active GFP-fusion protein (POI) could lose its function when it is sequestered to the 3×GFP aggregates. **(C)** Fluorescence microscopic images of WT (BY4741), *CDC20-GFP,* and *CDC28-GFP* strains under *3×GFP-op* and 3*×MOX-op*. Representative images are shown. **(D)** Quantification of cell size. For each strain, 100 randomly selected cells were analyzed. *: *p* < 0.05; N.S: *p* > 0.05; Tukey-Kramer method.

## Discussion

This study investigated how aggregate formation can be toxic or protective using 3×GFP as a model. In particular, we used 3×MOX, which is structurally similar but does not aggregate, as a control (Figure 1). This allowed us to separate the effects caused by protein overproduced from those caused by aggregates (Figure 2). Summarizing the results of the study, the mechanism of the large 3×GFP aggregate formation and exertion of toxicity can be explained as follows (Figure 8). 3×GFP-op induces aggregate formation, probably by partial misfolding during translation (Figure 1). The misfolded 3×GFP is ubiquitinated and degraded by the proteasome (Figure 3). This degradation is so active that it overloads the proteasome system, which causes growth defects During the formation process of the giant aggregates, smaller aggregates are surrounded and fused by the chaperone Hsp70, eventually transforming into a giant static mesh structure (Figure 5). In addition, during the process of aggregate enlargement, Hsp70 is trapped inside, triggering a protective HSR. This HSR constitutes a positive feedback loop that induces the expression of Hsp70 and leads to further enlargement of the aggregates. Although Hsp70 is trapped in the aggregates, it does not cause growth inhibition due to Hsp70 depletion, either because it is dynamic (Figure 4) or because the amount is sufficient (Figure 6). Therefore, the aggregates formed by 3×GFP-op themselves do not cause growth inhibition because they do not deplete essential proteins. On the contrary, it may be viewed as a necessary structure to capture Hsp70 to induce a productive heat shock response, as proposed by Klaips et al (Klaips et al. 2020). The positive correlation between the rate of aggregate formation and the decrease in growth inhibition also suggests that aggregate formation is cytoprotective. The formation of aggregates that are inaccessible to the proteasome might also play a role in reducing cytotoxicity by preventing the overloading of the proteasome.

The process of transformation of small aggregates into large dynamic Hsp70-capturing aggregates is considered to be equivalent to the observed transition between the two states of polyQ protein (Klaips et al. 2020). In normal yeast cells, polyQ protein forms rigid aggregates that do not trap enough Hsp70 to induce HSR. On the other hand, in cells overexpressing the co-chaperone Sis1, polyQ protein aggregates form a mesh structure that can trap dynamic Hsp70 (Figure 5). This causes a productive HSR. Time series analysis shows that the overproduced of 3×GFP initially results in the formation of small aggregates that co-localize with Hsp70. This structure is probably similar to that of polyQ protein aggregates when Sis1 is not overexpressed. Eventually, the small aggregates are surrounded by Hsp70 and fuse to form large aggregates that dynamically capture Hsp70 (Figure 4). This structure is probably similar to that of polyQ protein aggregates when Sis1 is overexpressed.

Protein overexpression results in growth inhibition (cytotoxicity) for a variety of reasons. We have categorized the mechanisms by which overexpression leads to toxicity into four categories: pathway modification, resource overload, stoichiometry imbalance, and promiscuous interactions (Moriya 2015). The formation of protein aggregates is thought to cause promiscuous interactions - loss of function by the non-specific trapping of essential proteins into the aggregates. 3×GFP aggregates trap essential Hsp70, but this does not appear to cause growth inhibition. On the other hand, overloading of the proteasome, i.e., degradation of resources, maybe the cause of growth inhibition. However, artificial trapping of essential cell cycle proteins into the 3×GFP aggregates caused cell cycle abnormalities (Figure 7), indicating that the toxicity of the aggregates is strongly context-dependent - what proteins can bind to them.

**Figure 8.**
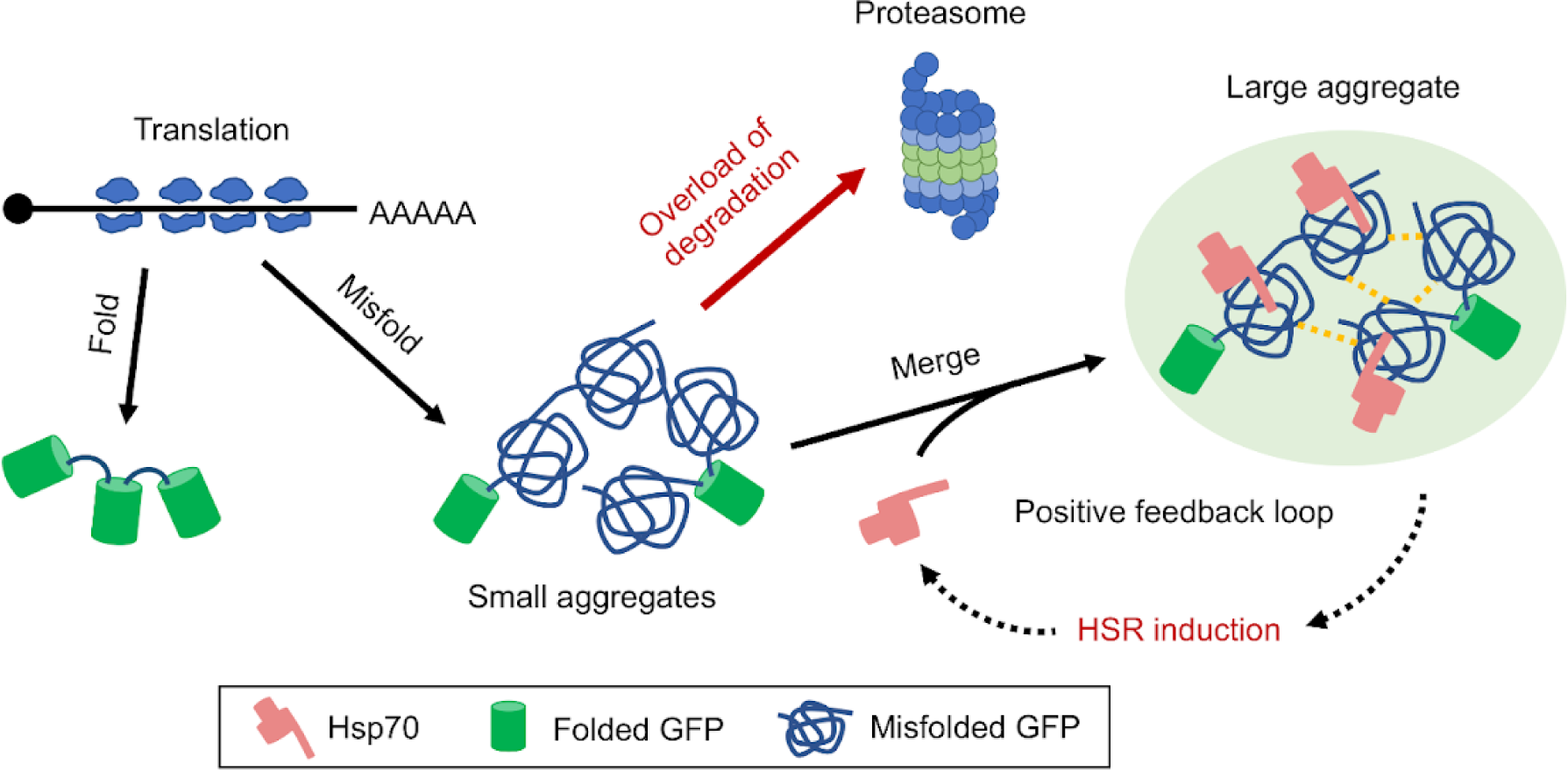
A hypothetical model of the effects caused by 3×GFP-op. The detail is explained in the main text.

## Materials and Methods

Strains, plasmids, assay kits, and reagents used in this study are listed in supplementary data 2.

### Strains and growth conditions

BY4741 (*MATa his3Δ1 leu2Δ0 met15Δ0 ura3Δ0*) were used as the budding yeast host strains (Brachmann et al. 1998). GFP collection, a temperature-sensitive mutant collection was created based on BY4741 (Z. Li et al. 2011; Huh et al. 2003). Cultivation and transformation of *S. cerevisiae* were performed as previously described (Burke, Dawson, and Stearns 2000). Synthetic complete (SC) medium without uracil (Ura) and leucine (Leu) as indicated was used for yeast culture. Cells were cultivated at 30°C otherwise noted.

### Plasmid construction

The plasmids were constructed by homologous recombination activity of yeast cells (Oldenburg et al. 1997), and their sequences were verified by DNA sequencing.

### Stress Condition

For the short-term induced stress of heat shock and H_2_O_2_, log phase cells cultured to OD660 = 0.8-1.0 were used. Heat shock stress was applied at 42°C for 10 min using a heat block. For H_2_O_2_, cells were left in SC–LU with H_2_O_2_ (0.3%) for 10 min. For AZC, yeast was liquid cultured in SC–LU with AZC (0.75 mM), and cells with OD660 = 0.8-1.0 were used.

### Measuring growth rate

Cellular growth was measured every 10 minutes using a microplate reader (680XR Biorad) for OD595. For temperature-sensitive proteasome mutants, an Infinite F200 microplate reader (Tecan) was utilized for temperature control. Maximum growth rate (MGR) was calculated as described previously (Moriya, Shimizu-Yoshida, and Kitano 2006).

### Microscopic observation

Log phase cells cultured to OD660 = 0.8-1.0 were observed with a fluorescence microscope (Leica DMI6000B). GFP fluorescence was observed with a GFP filter cube (Leica cat. # 11513899) and RFP fluorescence with an RFP filter cube (Leica cat. # 11513894) was used for observation. Image capture was performed using image acquisition software (Leica Application SuiteX).

### Image analysis

Cell recognition was conducted using YeastSpotter (Lu et al. 2019) to identify yeast cells from microscopic images (Figure S6A). Subsequent quantification of each cell’s size and brightness was performed using the raw fluorescence image data in CellProfiler (Version 4.2.0) (Stirling, Carpenter, and Cimini 2021). For cell size quantification in CellProfiler, the ‘MeasureObjectSizeShape’ module’s ‘Area’ parameter was employed, while brightness assessment was performed using the ‘Mean Intensity’ parameter from the ‘MeasureObjectIntensity’ module.

Aggregate recognition was primarily conducted through visual inspection. However, for screening involving spatial protein quality control-related genes, where analysis of over 100 images was necessary, aggregate segmentation was performed using Trainable Weka Segmentation within Fiji (Arganda-Carreras et al. 2017) (Figure S6A). The model was developed based on manually Segmented images (at least 100 cells), following the steps outlined in the official protocol. As a specific example, representative images are shown (Figure S6B).

### FRAP analysis

Samples were prepared as well as fluorescence microscopy observation. A confocal microscope (Olympus FV-3000) and a 100x oil lens were used. GFP was detected at ex/em = 488/540 and RFP was detected at ex/em = 561/670. Circular regions of a fixed size were bleached 3 frames after the start of the observation and then observed for 200 frames over 60 seconds. Bleaching was done using excitation light 488 nm. Image analysis was performed using Fiji (Schindelin et al. 2012). Fluorescence intensity data were normalized to 1 before light bleaching and 0 immediately after bleaching.

### Aggregate purification by his-tag

Cells were cultured in 25 ml of SC–LU medium overnight using 50 ml conical tubes. Cultured cells were washed with PBST with protease inhibitors (Thermo) and then crushed with a bead beater (TOMMY) (1 min x 4000 rpm) using glass beads. After crushing, cell extracts were centrifuged (20630 g x 10 min) and separated into soluble (sup) and insoluble fractions (ppt). Nickel beads (50µl) slurry was added The insoluble fraction was washed with 1 ml of PBST and centrifuged (20630 g x 5 min) 3-5 times. The insoluble fraction was then suspended in 1 ml of PBST and 50 μl of nickel carrier (his buffer kit, GE) was added. The samples were kept on ice for 30 minutes with occasional inverted mixing. At this stage, the binding of the aggregates to the nickel carrier could be confirmed by fluorescence microscopy. The carrier was then centrifuged at low speed (300 g x 1 min) and washed with PBST three times. 100 μl of PBST with 200 mM imidazole was added to the carrier and the aggregates were eluted. The carrier was then treated with 100 μl of NuPAGE LDS sample buffer (Thermo Fisher) at 70°C for 5 min, and subjected to SDS-PAGE.

### Total Protein extract

Cells overexpressing the target protein were cultured in SC–LU. Then, 1 ml of cells in the Log phase were collected (OD660 = 0.9-1.0). Cells were treated with 0.2 mol/L NaOH for 10 min (Kushnirov 2000) and then with 100 μl of NuPAGE LDS sample buffer (Thermo Fisher) for 5 min at 70·C.

### Protein analysis

For protein visualization, total proteins were labeled with Ezlabel FluoroNeo (ATTO) as described in the manufacturing protocol. SDS pages were then performed using 4-12% Gradient gel. Proteins were detected on a LAS-4000 (GE Healthcare) in SYBR-green fluorescence detection mode and Image Quant TL software.

Western blotting was performed to detect specific proteins. proteins separated by SDS-PAGE were transferred to a PVDF membrane (ThermoFisher). The membrane was then blocked with blocking buffer (PBST with 4% skim milk) for 1 hour, followed by incubation of the membrane with primary antibody diluted in PBST for 1 h. The membrane was then incubated with PBST containing peroxidase-conjugated secondary antibody (Nichirei bioscience) for 1. h. GFP was blocked with anti-GFP antibody (Roche) and ubiquitin was blocked with anti-ubiquitin was detected using an anti-GFP antibody (Roche) and ubiquitin was detected using an anti-ubiquitin antibody (Santa Cruz). The chemiluminescent image was acquired with a LAS-4000 image analyzer in chemiluminescence detection mode.

### RNAseq analysis

RNAseq analysis was performed essentially according to (Namba et al. 2022). Yeast overexpressing 3×GFP or 3×MOX was grown in SC–LU medium and collected in a log phase. For Vector controls, we used RNAseq data from (Namba et al. 2022) (accession number: GSE178244). RNA extraction was performed according to (Köhrer and Domdey 1991). Purified RNA was quality-checked by Multina (Shimazu). cDNA libraries were prepared using the TruSeq Stranded Total RNA kit (Illumina). Paired-end sequencing was performed using the Illumina NextSeq 550 (Illumina). Three biological duplications were analyzed for all strains. Sequences were checked for read quality by FastP (Chen et al. 2018) and aligned using Hisat2 (Kim et al. 2019). Aligned data were formatted into bam files by Samtools (H. Li et al. 2009) and quantified by StringTie (Pertea et al. 2015). Analysis of expression levels was performed by EdgeR (Robinson, McCarthy, and Smyth 2010). GO enrichment analysis was performed using the Gene Lists function on the SGD website (www.yeastgenome.org/).

## Acknowledgments

We would like to thank the members of the Moriya Laboratory (Okayama University) for their helpful discussions. We also thank Division of Instrumental Analysis, Okayama University for the FRAP measurements.

